# The effects of *CSTB* duplication on APP/amyloid-β pathology and cathepsin activity in a mouse model

**DOI:** 10.1101/2020.10.30.362004

**Authors:** Yixing Wu, Heather T. Whittaker, Suzanna Noy, Karen Cleverley, Veronique Brault, Yann Herault, Elizabeth M. C. Fisher, Frances K. Wiseman

## Abstract

People with Down syndrome (DS), caused by trisomy of chromosome 21 have a greatly increased risk of developing Alzheimer’s disease (AD). This is in part because of triplication of a chromosome 21 gene, *APP*. This gene encodes amyloid precursor protein, which is cleaved to form amyloid-β that accumulates in the brains of people who have AD. Recent experimental results demonstrate that a gene or genes on chromosome 21, other than *APP,* when triplicated significantly accelerate amyloid pathology in a transgenic mouse model of amyloid-β deposition. Multiple lines of evidence indicate that cysteine cathepsin activity influences APP cleavage and amyloid-β accumulation. Located on human chromosome 21 (Hsa21) is an endogenous inhibitor of cathepsin proteases, *CYSTATIN B* (*CSTB)* which is proposed to regulate cysteine cathepsin activity *in vivo*. Here we determined if three copies of the mouse gene *Cstb* is sufficient to modulate beta amyloid (Aβ) accumulation and cathepsin activity in a transgenic *APP* mouse model. Duplication of *Cstb* resulted in an increase in transcriptional and translational levels of *Cstb* in the mouse cortex but had no effect on the deposition of insoluble Aβ plaques or the levels of soluble or insoluble Aβ_42_, Aβ_40_, or Aβ_38_ in 6-month old mice. In addition, the increased CSTB did not alter the activity of cathepsin B enzyme in the cortex of 3-month old mice. These results indicate that the single-gene duplication of *Cstb* is insufficient to elicit a disease-modifying phenotype in the dupCstb x tgAPP mice, underscoring the complexity of the genetic basis of AD-DS and the importance of multiple gene interactions in disease.

## Introduction

Alzheimer’s Disease (AD) is the most common neurodegenerative disorder (1). Accumulation of amyloid-β (Aβ) plaques and formation of hyperphosphorylated tau tangles are pathological hallmarks of AD (2). People with Down’s syndrome (DS), a genetic disorder caused by chromosome 21 (Hsa21) trisomy develop the characteristic features of AD pathology by the age of 40 (3, 4) and by the age of 60 approximately 2/3 of individuals will have developed the clinical features of dementia (5). The amyloid precursor protein (APP) is encoded by a Hsa21 gene, *APP,* that is cleaved and processed to form Aβ which then aggregates to form plaques (6, 7). Duplication of *APP* is sufficient to cause early-onset AD (8, 9) and in the absence of three copies of *APP* people with DS do not develop early on set AD (10, 11). However, whether duplication of other Hsa21-located genes also contributes to the pathogenesis of AD in DS remains unknown.

In order to investigate the potential contribution of other Hsa21 genes to AD phenotypes, partial trisomy DS mouse models, which contain a segmental duplication of mouse chromosome regions that are syntenic to regions on Hsa21 have been generated (**Supplementary Figure 1**). Several segmental trisomy models for these regions have been generated and have been used to determine which combination of Hsa21 gene cause DS-associated phenotypes (12). Recently, by crossing a trisomic Hsa21 mouse model (Tc1), which has an extra copy of 75% Hsa21 genes but lacks an additional functional copy of *APP*, with an APP-amyloid deposition mouse model (J20-tgAPP) (13–15), we found that three copies of Hsa21 genes other than *APP* exacerbate Aβ deposition and cognitive deficits (16), indicating that other Hsa21 genes may also play important roles in the pathogenesis of DS-AD.

A potential candidate gene on Hsa21 is *CSTB*, which encodes cystatin B (CSTB), an endogenous inhibitor of cystine proteases (17). Human CSTB is an interacting partner of Aβ and colocalises with intracellular inclusions of Aβ in cultured cells (18). In the plaques of AD and Parkinsonism-dementia complex patients, protein levels of CSTB and cathepsin B are considerably increased (19). To date, several studies have provided evidence that cathepsin B influences amyloid pathology. However, the dominant mode of action is unclear. In secretory vesicles of neuronal chromaffin cells, cathepsin B inhibition disrupted the conversion of endogenous APP to Aβ (20). Treating an *APP* transgenic model (Tg(THY1-APP)2Somm) expressing the wildtype β-Secretase site of APP with cathepsin B inhibitors CA074Me or E64d leads to reduced Aβ levels and improved memory (21). These lines of evidence suggest a pro-amyloidogenic role of cathepsin B and a therapeutic potential for its endogenous inhibitor CSTB. However, knocking down cathepsin B in mice expressing familial AD-mutant human APP leads to increased levels of Aβ 1-42 and plaque deposition, indicating an anti-amyloidogenic role of cathepsin B (22). Knocking out *Cstb* by crossing the *Cstb*^*tm1Rm*^ (23) and an *APP* transgenic mouse model (TgCRND8) lowered cathepsin B activity (24), rescued autophagic lysosomal dysfunction and reduced Aβ aggregation (24), further indicating that CSTB may accelerate amyloid pathogenesis in the brain.

To determine if 3-copies of *CSTB* could influence the pathogenesis of AD in people with DS we crossed the J20 tgAPP mouse model of amyloid-β deposition to a mouse with a heterozygous duplication of the *Cstb* locus on Mmu10 (25). We found that duplication of *Cstb* increased transcriptional and translational levels of CSTB in the brain. However, this did not lead to changes in plaque deposition or Aβ levels. Duplication of *Cstb* did not change the activity of cathepsin B in the cortex at 3-months of age.

## Results

### *Cstb* mRNA and protein levels increased by duplication of *Cstb*

In order to verify the elevated transcript expression of *Cstb* in mice with a duplicated *Cstb* locus, the right cortices were prepared for qPCRs that compared the amount of *Cstb* mRNA to that of two different housekeeping genes (*Actb* and *Gapdh*). The results showed that duplication of *Cstb* in the mouse genome significantly increased the transcriptional level of *Cstb* (Figure 1A). The tgAPP had no effect on *Cstb* mRNA, nor was an interaction between tgAPP and dupCstb evident. Sex was not a significant factor (**Figure 1A**).

**Figure 1.**
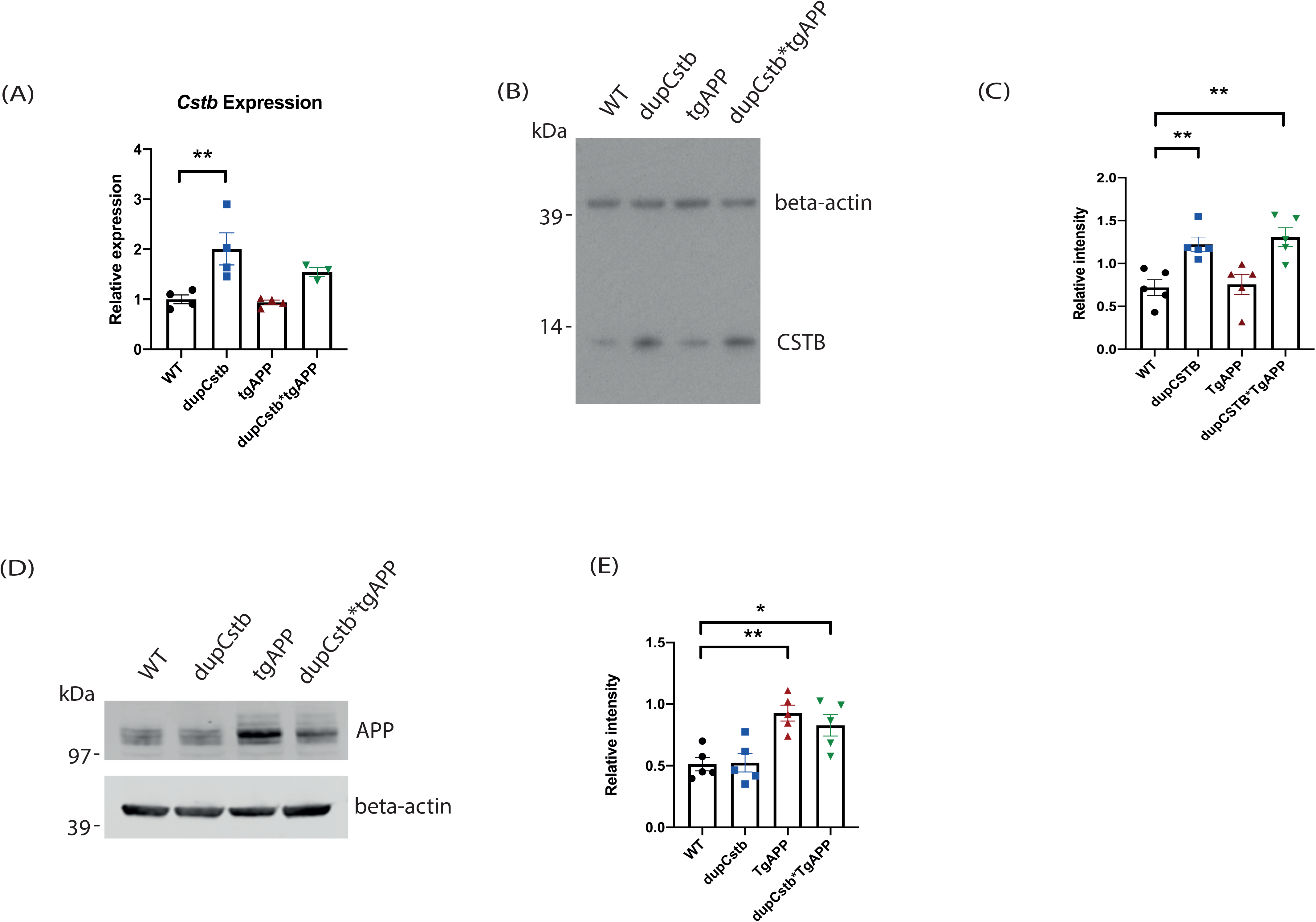
Transcriptional and translational levels of cortical CSTB in 3-month-old mice. (A) Levels of *Cstb* mRNA in the cortex of 3-month old mice. Data are relative to *Actb* and *Gapdh* housekeeping genes, and represented as mean expression levels for mice of each genotype ± SEM. Duplication of *Cstb* caused an increase in the relative levels of *Cstb* mRNA (F(1,7)=7.240, p=0.031). No effect of sex, tgAPP, or interaction between dupCstb and tgAPP was evident. (B) Representative western blot probed with anti-CSTB and anti-β-actin antibodies. (C) Protein band density was quantified using ImageJ, normalised to β-actin, and is shown as fold-change relative to wildtype levels (mean ± SEM), n=5 for each genotype (9 females and 11 males in total). (D) Representative western blot probed with anti-APP and anti-β-actin antibodies. (E) Protein band density was quantified using ImageJ, normalised to β-actin, and is shown as fold-change relative to wildtype levels (mean ± SEM), n=5 for each genotype (9 females and 11 males in total).

To determine whether the increased amount of mRNA translated to an increased amount of CSTB protein in dupCstb mice, cortical CSTB protein levels were measured by western blotting. The presence of a *Cstb* duplication caused an upregulation of CSTB protein (**Figure 1B**). The levels of CSTB were not changed due to tgAPP or sex, and there was no interaction observed between dupCstb and tgAPP. Mice containing dupCstb, including those in the dupCstb group and the dupCstb*tgAPP group, had approximately 2 times more CSTB protein than mice without dupCstb (**Figure 1C**). These data show that duplication of *Cstb* increases both the RNA and protein of CSTB in the cortex. Western blot was also used to confirm the overexpression of tgAPP (**Figure 1D**). tgAPP levels in the tgAPP group and the dupCstb*tgAPP group were significantly increased compared to mice without tgAPP (**Figure 1E**).

### *Cstb* duplication does not alter plaque deposition at 6 months of age in the cortex or hippocampus

We used the dupCstb*tgAPP mouse model to investigate if increased CSTB influences Aβ pathology at 6-months of age. The Aβ plaque load in the cortex or hippocampus was analysed using a 4G8 monoclonal antibody. Plaque deposition in both brain regions was apparent in mice with the tgAPP (**Figure 2A**). The 4G8 percentage coverage was calculated for two sections from every animal (**Figure 2B**). A significant effect of tgAPP was confirmed in the cortex, but there was no significant effect of a *Cstb* duplication or interaction between dupCstb and tgAPP. The results were also not significantly affected by the sex of the mouse.

**Figure 2.**
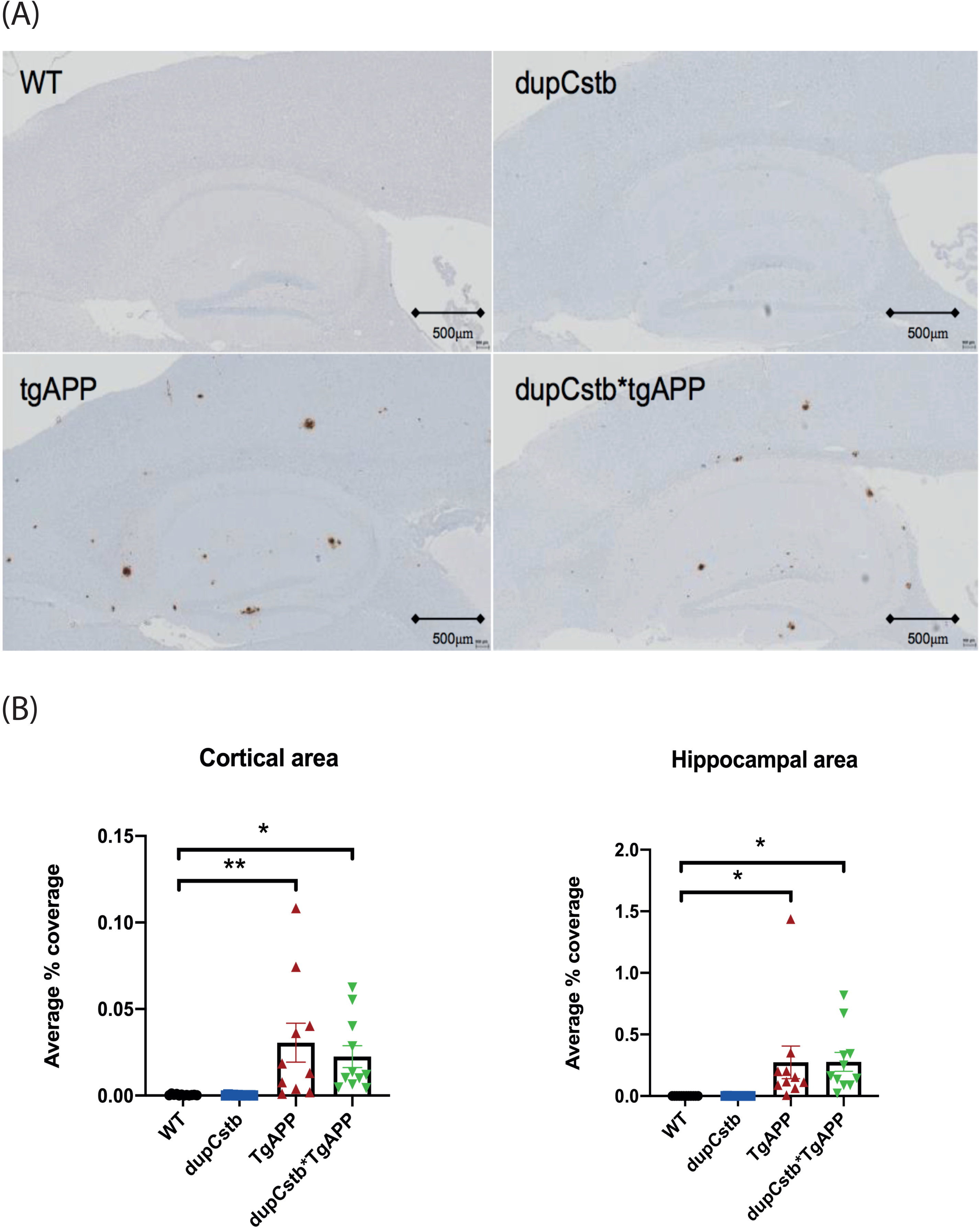
Aβ plaque deposition in the cortex and hippocampus of 6-month old mice. (A) Representative images of sagittal sections through the cortex and hippocampus. (B) Area stained with 4G8 antibody to Aβ was quantified as a percentage of the total area of that region, and group means ± SEM are presented. Mice with tgAPP had significantly more Aβ staining than those without, in the cortex (F(1,37)=19.503, p<0.001) and hippocampus (F(1,37)=14.466, p=0.001). There was no significant effect of Cstb duplication or sex, and no interaction between tgAPP and dupCstb. Data were analysed using a repeated measures ANOVA.

### *Cstb* duplication does not alter Aβ levels in the cortex at 6 months of age

We undertook a biochemical assay to probe the tissue for the quantity and solubility of Aβ_38_, Aβ_40_, and Aβ_42_ in the cortex from 6-month old mice. In both tgAPP and dupCstb*tgAPP groups, there was a differential representation of Aβ isoforms in the soluble and insoluble extracts (**Figure 3A-C**). We noted that in the Tris fraction Aβ_38_ was below the limit of detection in all but one sample, Aβ_38_ and Aβ_40_ predominating over Aβ_42_ in Triton-extracts (**Figure 3B**), and the reverse being true for Gnd-extracts (**Figure 3C**). Additionally, the values of Aβ in the Gnd-extracted fraction were a similar order of magnitude higher than in the Triton-soluble fraction for both genotypes measured. For example, the Aβ_42_ in Gnd extracts was on average 12,154 (± 3,111) pg/mg cortical protein in tgAPP and 9,567 (± 3,133) pg/mg in dupCstb*tgAPP mice, but in Triton-extracts Aβ_42_ was only 14.6 (± 3.19) pg/mg in tgAPP and 10.1 (± 1.45) pg/mg for dupCstb*tgAPP mice. The pattern displayed by both tgAPP and dupCstb*tgAPP groups is consistent with the expected Aβ biochemistry in the J20 mouse brain (26).

**Figure 3.**
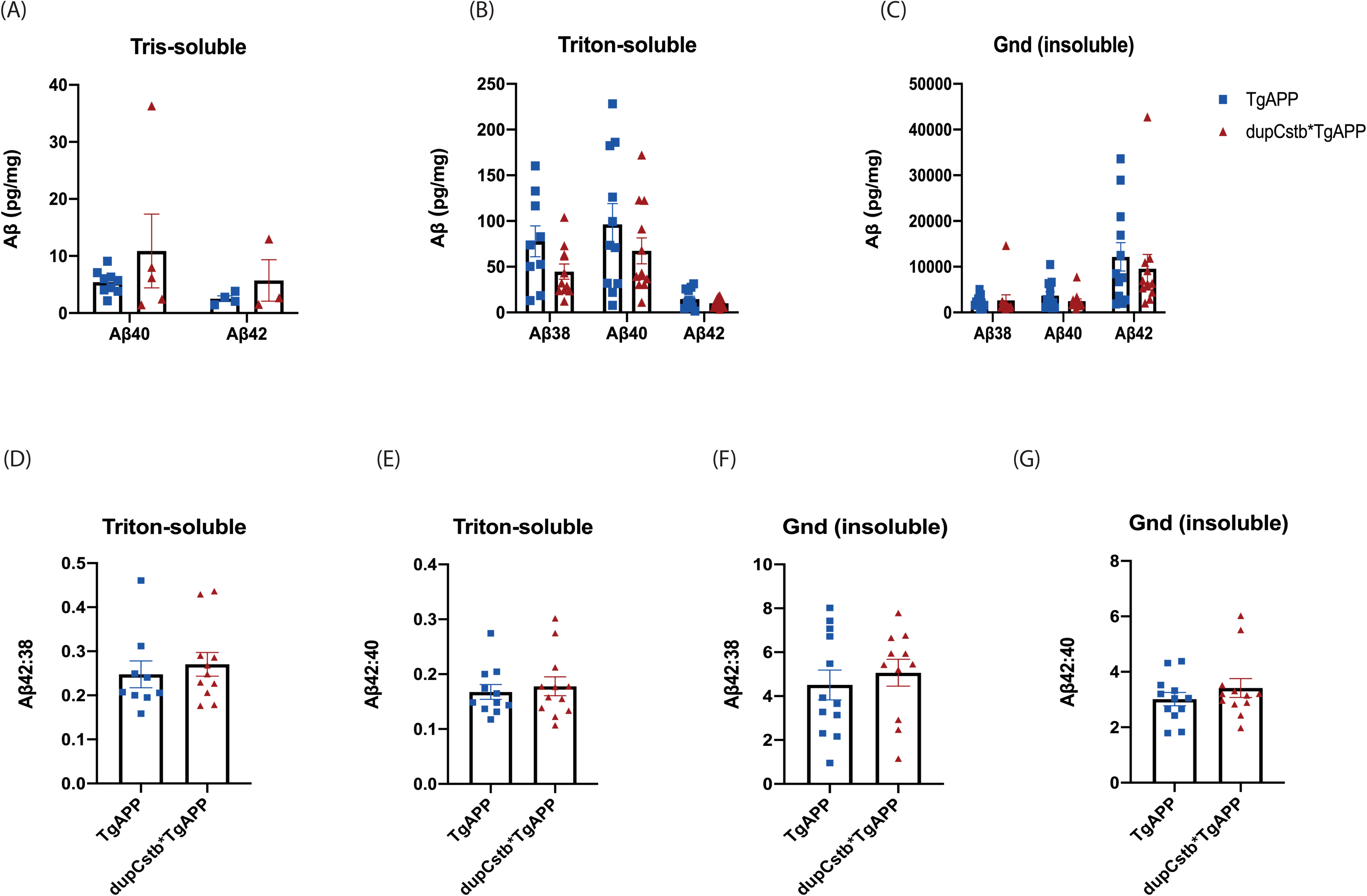
Cortical Aβ38, Aβ40, and Aβ42 in 6-month old mice, as measured by Meso Scale Discovery Assay. (A-C) Representation of group means for the Tris-, Triton-, and Gnd-soluble fractions. Samples below detection limit were recorded as ‘0’ in the graph. Values are in pg per mg total protein, ± SEM. (D-E) Ratio of Aβ42:Aβ38 and Aβ42:Aβ40 was calculated in the Triton-soluble fractions. (F-G) Ratio of Aβ42:Aβ38 and Aβ42:Aβ40 was calculated in the Gnd-soluble fractions. (A-G). No significant differences were found between tgAPP and dupCstb*tgAPP mice by univariate ANOVA.

To further investigate whether there were any changes in the Aβ peptide abundance due to dupCstb, the ratio of Aβ_42_:Aβ_38_ and Aβ_42_:Aβ_40_ were calculated for all mice carrying the tgAPP. The average ratio is presented for both tgAPP and dupCstb*tgAPP (**Figure 3D-G**), and shows that the presence of dupCstb did not have a statistically significant effect on this Aβ_42_:Aβ_38_ or Aβ_42_:Aβ_40_ metric in the cortex at 6 months of age. Together, these results indicate that *Cstb* duplication does not alter Aβ aggregation or Aβ ratios in the cortex at 6 months of age.

### *Cstb* duplication does not alter cathepsin B activity in the cortex at 3 months of age

To explore whether the increase in CSTB protein in dupCstb mouse brain corresponded to functional protease inhibition, a cathepsin B activity assay was conducted on cortex tissue from 3-month old mice. Activity was measured by the cleavage of a cathepsin B substrate to yield a fluorescence-emitting product. The mean rate of cathepsin B activity in each cohort of littermates as a percentage of the WT mean rate was calculated (**Figure 4**). Univariate ANOVA revealed no significant effect of dupCstb, tgAPP, or sex, and no interaction between these factors, suggesting that upregulation of *Cstb* has little impact on cathepsin B activity regulation in the cortex at 3 months of age.

**Figure 4.**
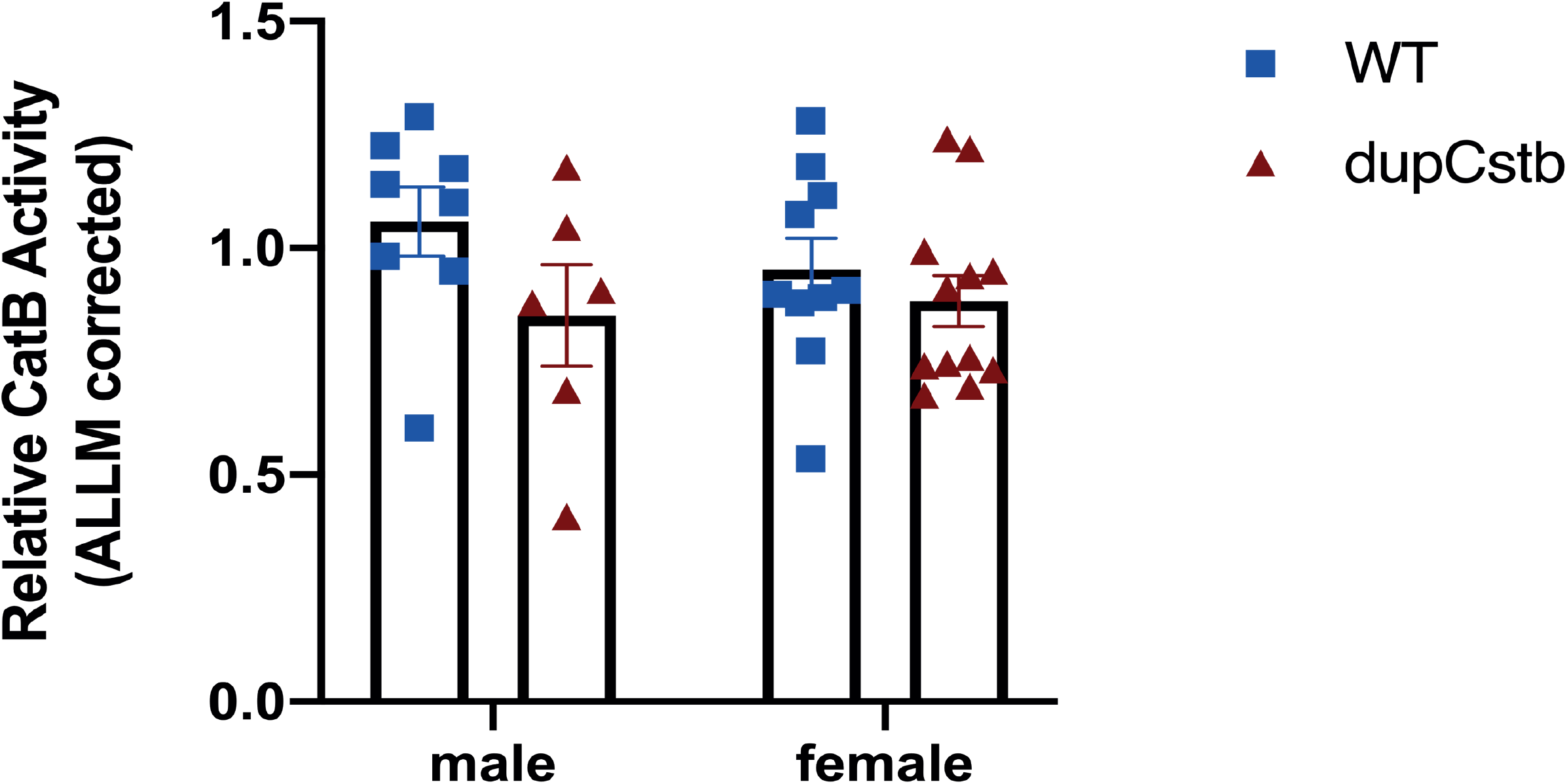
Activity of cathepsin B in cortical lysates of 3-month-old mice. Enzyme activity, as the gain in fluorescence during the linear portion of the reaction curve was calculated relative to the wildtype mean. These data, graphed as group means ± SEM, n=18 for each genotype (22 females and 14 males in total), revealed no statistically significant effects of dupCstb or sex (univariate ANOVA).

## Discussion

In this study, we showed that despite a heterozygous duplication of the *Cstb* gene leading to an upregulation of CSTB protein in the mouse brain, this had little impact on the degree of amyloid-β plaque deposition in the J20 tgAPP model at 6 months-of-age. In contrast, by crossing the *Cstb* knockout (*Cstb*^*tm1Rm*^) mouse with the tgAPP TgCRND8 model (27), plaque load was significantly reduced at 6-months of age (24), indicating a potential therapeutic importance of *CSTB* knockout. Our data on the null effect of *Cstb* duplication are not necessarily conflicting with the result using the *Cstb*^*tm1Rm*^ mouse model, given that loss of and gain of function often have differing affects.

Transgene overexpression in tgAPP mouse models is frequently criticised as being unrepresentative of human fAD and especially sAD (28, 29). Although some of these mouse models have proven to be valuable reductionist platforms for experimental manipulation, the mutant human gene is expressed at far higher levels than is physiologically relevant to the disease. In this study the high level of APP and robust production of Aβ may not be modifiable by a single additional copy of the *Cstb* gene. Similarly, the results of specific gene ‘triplication’ studies should be extrapolated with caution to trisomy of Hsa21, in which transcriptional dysregulation has been documented and may be an effect of aneuploidy in general (30, 31).

Interestingly, localised γ-secretase activity that is selective for lysosomal substrates has been shown to generate an intracellular pool of Aβ_42_ (32) and this pool may be affected in a model of lysosomal dysregulation. CSTB is localised in the nucleus, cytosol and lysosome (33). Although it is an endogenous inhibitor of cystine cathepsins (34), the subcellular distribution of CSTB is important in determining and maintaining its regulatory role. Whether the increased level of CSTB in our model is lysosome associated is unknown. In this study, the lack of significant changes in soluble Aβ suggests that there are not any changes to intracellular Aβ but further experimentation is required to verify this. We note that early intracellular depositions of amyloid-β is a prominent feature of the earliest stages of AD-DS (35–38).

Cathepsin B enzymatic activity was found to be elevated in brains of Cstb knockout mice (24). However, our study was unable to demonstrate the inverse at 3 months of age. Aging may modulate the effect of *Cstb* duplication on cathepsin B enzymatic activity, indeed previous studies have suggested the aging exacerbates the effect of trisomy (39). A further study would be required to investigate this.

Overall, our study reveals that duplication of the *Cstb* gene alone is unlikely to modify APP/Aβ pathogenesis in the mouse model used. These results prompted the question of whether having 3-copies of *Cstb* are sufficient to elicit a functional consequence on cathepsin activity, or whether the lack of change was due to a compensatory downstream mechanism. In order to conclude that 3-copies of *CSTB* are necessary for exacerbating pathology in the context of Hsa21 trisomy, the Tc1*J20 mouse model would need to be crossed with a *Cstb* knockout mouse.

## Material and methods

### Mouse breeding and husbandry

Mice with a duplication of *Cstb* (CSTB5HP mice, EMMA stock code EM:04463, MGI:5828767, named here dupCstb) were re-derived from embryos and maintained in a colony by mating dupCstb males to C57BL/6J females. B6.Cg-Zbtb20^Tg(PDGFB-APPSwInd)20Lms^/2Mmjax (MGI:3057148, named here J20) were obtained from a colony maintained by mating J20 mice (JAX stock code 0006293) to C57BL/6J mice. DupCstb females were mated with J20 males to produce the dupCstb x J20 colony. This colony produced mice with four genotypes referred to as: wildtype (WT), dupCstb, tgAPP, and dupCstb*tgAPP.

Genotyping of mice was outsourced to Transnetyx. qPCR was performed to check for any reduction in human *APP* copy number in the tgAPP mice, to exclude from analysis any with a copy number dropped by at least 40% as compared to a J20 positive control DNA, from the Jackson laboratory, Bar Harbor, Maine.

The mice involved in this study were housed in controlled conditions in accordance with Medical Research Council guidance (*Responsibility in the Use of Animals for Medical Research*, 1993), and experiments were approved by the Local Ethical Review panel and conducted under License from the UK Home Office. Cage groups and genotypes were semi-randomised, with a minimum of two mice to a cage and litters; group weaned with members of the same sex. Mouse houses, bedding and wood chips, and continual access to water were available to all mice, with RM1 and RM3 chow (Special Diet Services, UK) provided to breeding and stock mice, respectively. Cages were individually ventilated in a specific-pathogen-free facility. Mice were euthanised by exposure to a rising concentration of CO_2_ gas, according to the Animals (Scientific Procedures) Act issued in the United Kingdom in 1986.

### Histology

Immediately following euthanasia, the brain was removed, dissected sagittally along the midline and the left hemisphere immerse fixed in 10% buffered formal saline for 48-72 hours (Pioneer Research Chemicals, UK). The fixed tissue was embedded in paraffin wax using an Automated Vacuum Tissue Processor (Leica ASP 300S, Germany) and a series of 4μm sections with the hippocampal formation were cut and mounted onto Superfrost plus glass slides. For amyloid-β immunostaining sections were dewaxed, rehydrated through an alcohol series to water, pre-treated with 80% formic acid for 8mins followed by washing in distilled water for 5mins. The sections were stained for amyloid-β using the Ventana Discovery XT automated stainer, where further pre-treatment (mild CC1 - 30 minutes of EDTA Boric Acid Buffer, pH 9.0), and blocking (8mins (Superblock, Medite, #88-4101-00), were performed prior to primary antibody incubation (12hrs, biotinylated mouse monoclonal antibody, Sigma-Aldrich SIG-39240 Beta-Amyloid - 4G8) at 2μg/ml (antibody diluent, Roche, Switzerland). The staining was completed with the Ventana XT DABMap kit and a haematoxylin counterstain, followed by dehydration and permanent mounting with DPX. All images were acquired using a Leica SCN400F slide scanner analysed using Definiens Tissue Studio and Developer software, with regions of interest manually outlined with reference to a mouse brain atlas (40). A single operator performed segmentation of all the images, which the software then processed to quantify the area of DAB staining.

### Quantitative reverse transcriptase PCR (qRT-PCR) for *Cstb*

Cortical RNA was extracted as per the miRNeasy Mini Kit protocol (QIAGEN, January 2011), and cDNA was then generated using a QuantiTect Reverse Transcription Kit (QIAGEN), including genomic DNA (gDNA) elimination. TaqMan® quantification using a VIC-*Actb* probe (#4331182) or VIC-*Gapdh* probe (#4331182) and FAM-*Cstb* probe (#451372) was undertaken using a 7500 Fast Real Time PCR System (Applied Biosystems. A 2-fold serial dilution sample of WT mouse cDNA was used as a standard.

### Western blotting

For analysis of protein abundance, cortex was dissected under ice-cold PBS before snap freezing. Samples were then homogenized in RIPA Buffer (150 mM sodium chloride, 50 mM Tris, 1% NP-40, 0.5% sodium deoxycholate, 0.1% sodium dodecyl sulphate) plus complete protease inhibitors (Calbiochem) by mechanical disruption. Total protein content was determined by Bradford assay. Samples from individual animals were run separately and were not pooled.

Cortical homogenates were denatured in NuPAGE**®**LDS Sample Buffer (Life Technologies, USA) and 2μl 2% β-mercaptoethanol (Sigma-Aldrich) at 100°C for 5 minutes and separated by SDS-polyacrylamide gel electrophoresis on a NuPAGE**®**Novex**®**4-12% Bis-Tris at 200V for 30 minutes. The proteins in the gel were transferred to a nitrocellulose membrane by Trans-Blot® Turbo™ Transfer System (Bio-Rad Laboratories) at 25V, 2.5A, for 15 minutes. The membranes were blocked with 5% (w/v) milk in PBST (PBS with 0.05% Tween-20 (Sigma-Aldrich)) or Intercept® (PBS) Blocking Buffer (Selected P/N: 927-70001, LI-COR, USA) for 1 hour at room temperature. The membranes were then incubated in primary antibodies overnight at 4°C, an anti-β-actin mouse monoclonal antibody (Sigma-Aldrich, #A5441, 1:5,000) or a rabbit anti-APP antibody (Sigma, USA, #A8717, 1:4,000) or an anti-Cystatin B rat monoclonal antibody (Novus Biologicals, USA, #227818, 1:2,000). Followed-by Goat anti-Rat IgG (H+L) Secondary Antibody HRP (Thermo Scientific, USA, # 31470 1:5,000), Goat anti-Rabbit IgG H&L (IRDye® 800CW, Abcam, ab216773, 1:10,000) or Goat anti-Mouse IgG H&L (IRDye® 800CW, Abcam, ab216772, 1:10,000) for 1 hour at room temperature. Once antibody probing was complete, for membranes labelled with secondary antibody HRP, SuperSignal™ West Pico Substrate (Thermo Scientific) was used for chemiluminescent detection of bound proteins, the membranes were exposed to Hyperfilm ECL (GE Healthcare Life Sciences, USA, #10607665) for 30 seconds, and visualised on a ChemiDoc MP Imaging System (Bio-Rad). Membranes labelled with IRDye® 800CW were visualised using Odyssey CLx Infrared Imaging System (LI-COR). The density of protein bands was analysed with Image-J. The calculated density of the band corresponding to CSTB was divided by the density of the β-actin band. To normalise the data to each blot and facilitate comparison of data between blots, the average of two WT relative CSTB values on each blot was taken, and the density values of all lanes on that blot was divided by the WT average.

### Biochemical Fractionation and Meso Scale Discovery Assay

Weighed cortex was homogenised in Tris-buffered saline (TBS) (50 mM Tris-HCl pH 8.0) plus complete protease and phosphatase inhibitors (Calbiochem) before centrifugation at 175,000 × g for 30 minutes at 4°C. The supernatant was removed, snap frozen, and stored at −80°C as the Tris-soluble fraction. The remaining pellet was re-suspended by homogenising in ice cold 1% Triton-X (Sigma-Aldrich) in TBS (pH 8.0) and centrifuged at 175,000 × g for 30 minutes at 4°C. The resultant supernatant was removed, snap frozen, and stored at −80°C as the Triton-soluble fraction. The pellet was re-homogenised in 500μl ice cold 5M Guanidine (Gnd) HCl (Sigma-Aldrich) in TBS (pH 8.0). The final volume was brought to 8x the half-cortex weight with 5M Gnd-TBS, and left rocking overnight at 4°C to fully re-suspend the sample. This fraction was then snap frozen and stored at −80°C as the Gnd-soluble fraction. Each biochemical fraction was individually assayed for levels of Aβ_38_, Aβ_40_, and Aβ_42_ using an MSD**®**MULTI-SPOT Human (6E10) Abeta Triplex Assay (MesoScale Discovery, USA, #K15148E-2) in duplicate. The protocol was carried out as per manufacturer instructions, using reagents provided with the kit; samples were diluted 1:1 (for Tris- and Triton-soluble fractions) or 1:20 (Gnd-soluble fraction) in Diluent 35. Peptide abundance was normalised to wet weight of cortex.

### Cathepsin B activity assay

Cathepsin B activity was assayed in the cortex of 3-month old mice using a fluorometric kit (Abcam, #ab65300). Tissue was homogenised in lysis buffer then incubated for 30 minutes on ice, then prior to centrifugation at 15,000 x g for 5 minutes at 4°C. The supernatant was removed and transferred to a clean tube, on ice, and the protein concentration was determined by a Bradford assay. For each sample, a volume of supernatant containing 50μg protein used for each reaction with cathepsin B Substrate (RR-amino-4-trifluoromethyl coumarin (AFC)).

To control for non-specific cleavage, negative controls treated with inhibitor ALLM (Abcam, ab141446), were run for every sample. The reaction was incubated at 37°C and the fluorescent output (excitation/emission = 400/505nm) was recorded every 90 seconds for 30 cycles. The linear range of the reaction was determined and the relative cathepsin B activity in the sample calculated by taking the average fluorescent output for each sample minus that in the inhibited. Values are expressed as a percentage of WT average.

### Experimental Design and Statistical analysis

All experiments and data analysis were undertaken blind to genotype and sex of the mouse. Graphs were plotted using Prism8 (GraphPad). All statistical analysis was carried out using SPSS Statistics 22 (IBM). Univariate or repeated measures analysis of variance (ANOVA) analyses were used to assess the contribution of tgAPP, dupCstb, and sex of the mouse, to the outcome variable, as well as any interaction between these factors.

## Acknowledgement

Y. W. is funded by an Alzheimer’s Research UK Senior Research Fellowship held by F.K.W (ARUK-SRF2018A-001). F.K.W. is also supported by the UK Dementia Research Institute which receives its funding from DRI Ltd, funded by the UK Medical Research Council, Alzheimer’s Society and Alzheimer’s Research UK. F.K.W. and E.M.C.F. also received funding that contributed to the work in this paper from the MRC via CoEN award MR/S005145/1. E.M.C.F. received funding from a Wellcome Trust Strategic Award (grant number: 098330/Z/12/Z) awarded to The London Down Syndrome (LonDownS) Consortium (E.M.C.F) and a Wellcome Trust Joint Senior Investigators Award (grant numbers: 098328, 098327).

We thank Dr. Amanda Heslegrave (UCL-DRI) for assistance with this project.

F.K.W. has undertaken consultancy for Elkington and Fife Patent Lawyers unrelated to the work in the manuscript.

---

**Supplementary figure 1.**
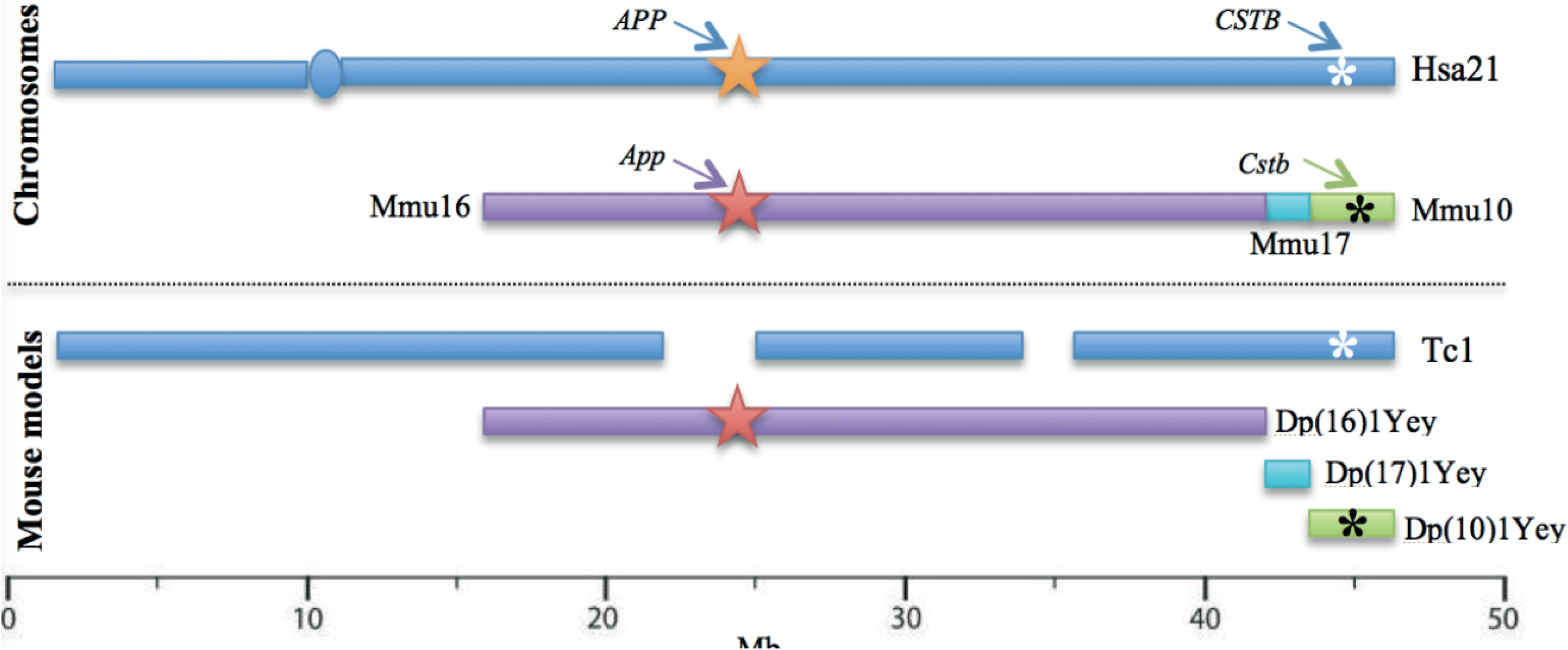
Schematic diagram of mouse models trisomic for Hsa21 orthologous genes adapted from Choong et al. (2015). The regions of Mmu16, 17, and 10 are aligned with their corresponding regions on the long arm of Hsa21, along a megabase pair (Mb) scale. The Tc1 mouse model is represented with breakpoints excluding the Hsa21 genes that are not functionally expressed. The approximate position of the human and mouse APP and CSTB genes are indicated with arrows.

## Notes

### Competing Interest Statement

The authors have declared no competing interest.

